# Metabolic Sensing of Extra-cytoplasmic Copper Availability via Translational Control by a Nascent Exported Protein

**DOI:** 10.1101/2022.11.11.516240

**Authors:** Yavuz Öztürk, Andreea Andrei, Crysten E. Blaby-Haas, Noel Daum, Fevzi Daldal, Hans-Georg Koch

## Abstract

Metabolic sensing is a crucial prerequisite for cells to adjust their physiology to rapidly changing environments. In bacteria, the response to intra- and extra-cellular ligands is primarily controlled by transcriptional regulators, which activate or repress gene expression to ensure metabolic acclimation. Translational control, such as ribosomal stalling can also contribute to cellular acclimation and has been shown to mediate responses to changing intracellular molecules. In the current study, we demonstrate that co-translational export of the protein CutF regulates translation of the down-stream *cutO*-encoded multi-copper oxidase CutO in response to extracellular copper (Cu). Our data show that CutF, acting as a Cu sensor, is co-translationally exported by the signal recognition particle pathway. Binding of Cu to the periplasmically exposed Cu-binding motif of CutF delays its co-translational export via its C-terminal ribosome stalling-like motif. This allows the unfolding of an mRNA stem-loop sequence that shields the ribosome-binding site of *cutO,* which favors its subsequent translation. Bioinformatic analyses reveal that CutF-like proteins are widely distributed in bacteria and often, are located upstream of genes involved in transition metal homeostasis. Our overall findings illustrate a highly conserved control mechanism using co-translational export of a protein acting as a sensor to integrate the changing availability of extracellular nutrients into metabolic acclimation.

**Importance:** Metabolite sensing is a fundamental biological process, and the perception of dynamic changes in the extracellular environment is of paramount importance for the survival of organisms. Bacteria usually adjust their metabolism to changing environments by transcriptional regulation. Here, we describe an alternative translational mechanism that controls the bacterial response to the presence of copper, a toxic micronutrient. This mechanism involves a co-translationally secreted protein that, in the presence of copper, undergoes a process resembling ribosomal stalling. This allows the unfolding of a downstream mRNA stem-loop and enables translation of the adjacent Cu-detoxifying multicopper oxidase. Bioinformatic analyses reveal that such proteins are widespread, suggesting that metabolic sensing using ribosome-arrested nascent secreted proteins acting as sensors may be a common strategy for integrating environmental signals into metabolic adaptation.

## Introduction

Living cells have evolved complex sensing and signaling systems for ensuring survival in rapidly changing environments. This is particularly important for single-cell organisms, such as bacteria. They engage multiple sensing systems in parallel, including transcriptional regulators (1), two-component systems (2) and chemotactic responses (3). Metabolite sensing by these systems guarantees sufficient nutrient supply while simultaneously helping to evade potentially toxic compounds. However, the situation is more complex for micronutrients, which are essential for cell metabolism, but could be toxic even at low concentrations. This is exemplified by the multifaceted bacterial response to variations in the concentration of copper (Cu) (4).

Cu is an essential micronutrient used as a cofactor by cuproenzymes, such as cytochrome oxidases (Cox) or nitrous oxide reductases (4–6). However, free Cu is also toxic due to its redox properties, as it facilitates the formation of reactive oxygen species, interferes with thiol groups in proteins and damages various metalloproteins (6, 7). Consequently, no free Cu is detectable in the bacterial cytoplasm (8), with Cu homeostasis achieved by an intricate network of Cu transporters and Cu chaperones. This ensures sufficient Cu supply for cuproprotein biogenesis while preventing accumulation of free Cu (4, 9–12). Periplasmic multi-copper oxidases provide a further defense line against Cu toxicity by converting Cu(I) to the less toxic Cu(II) (13–15).

Transcriptional activation of genes encoding Cu-exporting proteins and Cu-chaperones represent the main bacterial response to increased Cu concentrations (16–18). MerR-like transcriptional activators, such as CueR, sense Cu in the cytoplasm and regulate the production of the cytosolic Cu chaperone CopZ, the Cu-exporting P_1B_-type ATPase CopA, and the multi-copper oxidase CutO (19–21). Transcriptional repressors also regulate Cu-dependent gene expression. At low cytoplasmic Cu concentrations, repressors such as CsoR, CopY, or ArsR prevent transcription of genes encoding Cu-exporting P_1B_-type ATPases and Cu chaperones. On the other hand, periplasmic Cu concentrations are primarily sensed by two-component systems, such as CusRS or CopRS (22).

In addition to transcriptional control, some Cu-response proteins are post-transcriptionally regulated as well. One example is the proteolysis of CueR by the AAA^+^ proteases Lon, ClpXP and ClpAP (23). Another example is the multi-copper oxidase CutO in *Rhodobacter capsulatus* (15, 24). In response to Cu, production of CutO is regulated by an mRNA stem-loop sequence (SL) located between *cutF* and *cutO* and strictly requires the presence of an intact *cutF* in the tricistronic *cutFOG* operon (**Fig. 1A**). Although *cutF* can be translated into protein *in vitro*, CutF is not detectable *in vivo* by immune detection or mass spectrometry (15, 25). Moreover, genetic complementation of a Δ*cutF* mutant is only possible when the *cutFOG* operon is kept intact, indicating that *cutF* must be located immediately upstream of *cutO* to execute its function. Bioinformatic searches identified CutF as a member of the DUF2946-like family of proteins, which are abundant in Pseudomonadota and are often encoded upstream of cuproenzymes, periplasmic Cu chaperones, and Cu transporters (15).

**Figure 1.**
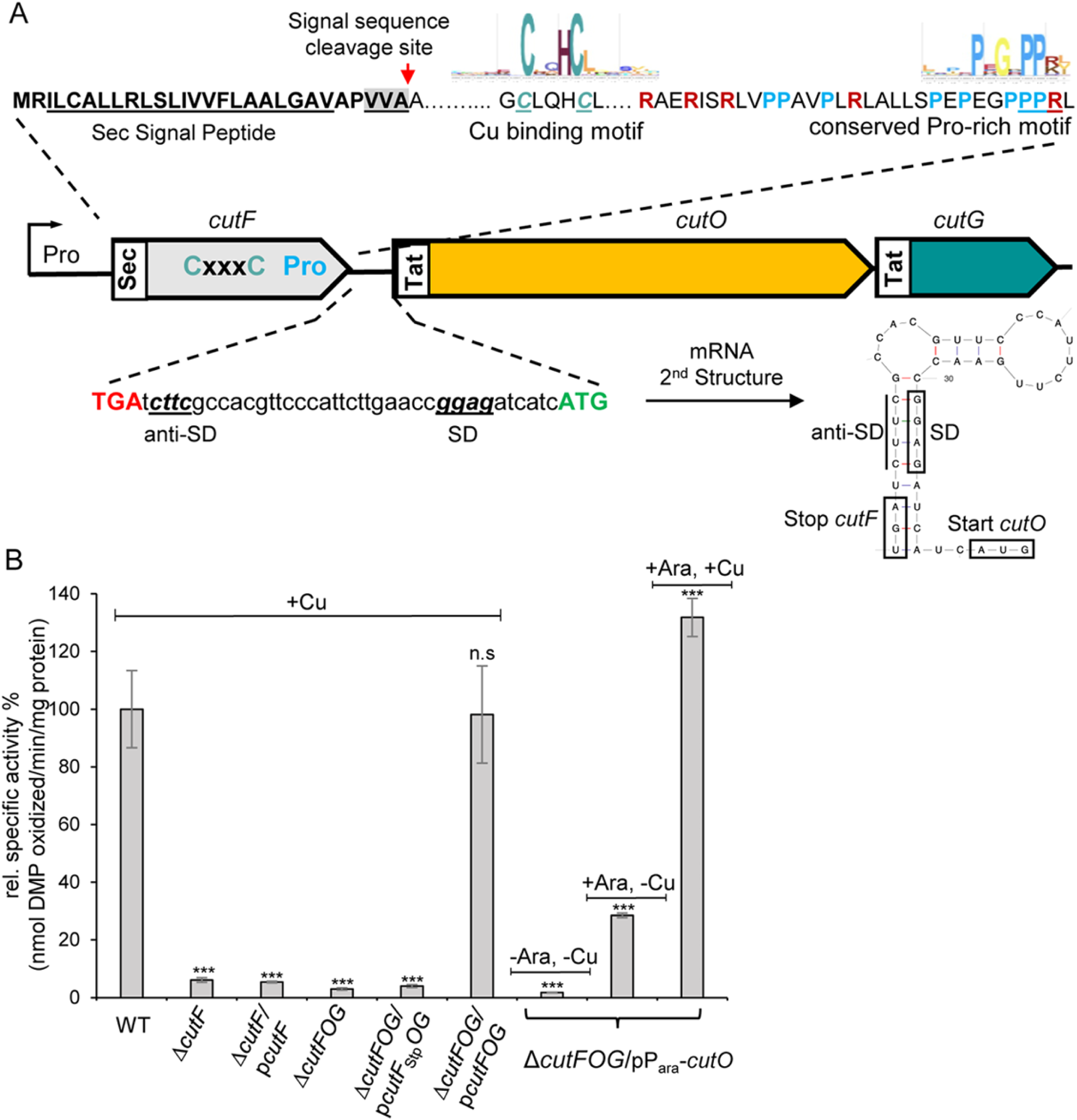
The structural integrity of the *cutFOG* operon is essential for CutO production. (**A**) Genetic organization of the *cutFOG* operon in *Rhodobacter capsulatus*. The *cutFOG* operon encodes CutF, a predicted protein of unknown function, the multi-copper oxidase CutO and the copper chaperone CutG. CutF has a cleavable Sec signal sequence (Sec) and CutO and CutG contain a twin-arginine signal sequence (Tat). The putative Cu-binding motif (CxxxC) and a proline-rich C-terminus of CutF are indicated. The intergenic *cutF-cutG* region forms a stem-loop structure, which shields the Shine-Dalgarno sequence (SD) of *cutO*. (**B**) CutO activity of different *R. capsulatus* strains that either lack *cutF* (Δ*cutF*) or the entire *cutFOG* operon (Δ*cutFOG*). p*cutF* contains an N-terminally Flag-tagged copy of *cutF* under the control of its native promoter in the low-copy plasmid pRK415, and similarly, p*cutFOG* encodes epitope-tagged versions of the entire *cutFOG* operon with an N-terminal Flag-tag, a C-terminal Flag-tag, and a C-terminal Myc-His-tag, respectively. The plasmid p*cutF*_Stp_*OG* is a derivative of p*cutFOG* with a substitution of the *cutF* start codon with a stop codon. The plasmid pP_ara_-*cutO* contains the C-terminally Flag-tagged CutO under arabinose promoter control. CutO activities were determined by the 2,6-DMP assay, using the periplasmic fractions (50µg total protein) of appropriate strains after growth on MPYE enriched medium. When indicated, MPYE medium was supplemented with 10 µM CuSO_4_ and 0.5 % arabinose. The activity of wild-type (WT) was set to 100%, and the relative activities of the indicated strains were calculated. Three independent experiments and in each case three technical replicates were performed, and the error bars reflect the standard deviation (n=9). Statistical analyses were performed with the Satterthwaite corrected two-sided Student t-test, using the activity of the WT as a reference. (*) refers to p-values ≤0.05; (**) to p-values ≤0.01, (***) to p-values ≤0.001 and n.s. to not significant.

Despite their abundance, the function of CutF-like proteins is unknown. Our findings show that co-translational export of the putative Cu-binding protein CutF is delayed by a Cu-induced translational pausing-like process, causing unfolding of a downstream mRNA SL sequence which allows translation of the adjacent *cutO*. Bioinformatic analyses reveal that such metabolic sensing mechanisms may represent a more common and largely unexplored regulatory principle in bacteria.

## Results

### Structural integrity of *cutFOG* operon and translation of *cutF* is required for Cu-dependent CutO production

CutF is essential for CutO production (15) with a Δ*cutF* strain showing the same low CutO activity as a Δ*cutFOG* strain (**Fig. 1B**). While the Δ*cutFOG* strain is fully complemented by ectopic expression of the *cutFOG* operon, the lack of CutO activity and the associated Cu-sensitivity of a non-polar Δ*cutF* mutant cannot be complemented *in trans* by a plasmid expressing solely *cutF* (p*cutF*) (**Fig. 1B**, **Fig. S1A**). This indicates that the *cutF* product can execute its function only when in *cis* to *cutO*. Since CutF is not detectable in cells (15) 25), whether *cutF* is translated *in vivo* was probed by replacing its predicted ATG start codon with a stop codon in a plasmid containing the intact *cutFOG* operon (p*cutF_Stp_OG*). This construct failed to restore CutO activity, whereas the same plasmid containing *cutF* with its start codon (p*cutFOG*) allows for wild-type CutO activity and the associated Cu-tolerance **(Fig. 1B, Fig. S1B)**. Thus, *cutF* translation *in vivo* is needed to support Cu-dependent CutO production.

Analysis of the *cutFOG* transcript shows that *cutO* is preceded by a stem-loop (SL) structure that shields the predicted *cutO* Shine-Dalgarno sequence (SD) (24) (**Fig. 1A**). Unfolding of this region likely required for CutO synthesis. Whether CutF is required for SL unfolding was addressed by ectopic expression of *cutO* under the control of an arabinose-controlled promoter in the Δ*cutFOG* strain in the absence of SL and CutF (pP_ara_-*cutO*). CutO activity was detectable upon addition of arabinose, and this activity increased by Cu supplementation (**Fig. 1B**). Immune-detection showed that Cu barely affects the steady-state amount of Flag-tagged CutO **(Fig. S1C)**, indicating that Cu addition has no major effect on CutO levels, but rather enhances CutO enzymatic activity, which has been also observed before (15) Thus, CutO can be readily produced in the absence of both CutF and SL, when expressed from a *cutO* gene in trans.

Whether CutF is required for Cu- and SL-dependent production of CutO was tested using two constructs. Both lacked *cutF*, but only one contained SL **(Fig. S2A)**. In the presence of SL and absence of *cutF* (p*cut(+SL)OG*), no significant CutO activity was observed, whereas in the absence of both SL and *cutF* (p*cut(-SL)OG*) some CutO activity (**Fig. 2A**), and the associated Cu-tolerance **(Fig. S1D),** were detectable. Considering that the SL could shield the SD sequence upstream of *cutO*, a third construct was tested in which the anti-SD sequence was mutated (24) **(Fig. S2A)**. The CutO activity in Cu-supplemented cells containing this mutated SL (*SL_m_*) and lacking *cutF* (*cut(SLm)OG*) was higher than the activity observed in the absence of SL. This suggested that eliminating the *cutF-SL* region also interferes with the stability of the *cutOG* mRNA, as seen earlier (15, 24). Further supporting this hypothesis, CutO activity increased almost ten-fold in cell extracts expressing *cutF* and *SLm* together (*cutF_SLm_OG*) (**Fig. 2A**), and immune detection showed that CutO levels were strongly increased in cell extracts **(Fig. S2B,** *cutF_SLm_OG*). This showed that CutF is required for maximum Cu-dependent CutO production only in the presence of an intact *cutF-SL* region, and not in the absence of the SL

**Figure 2.**
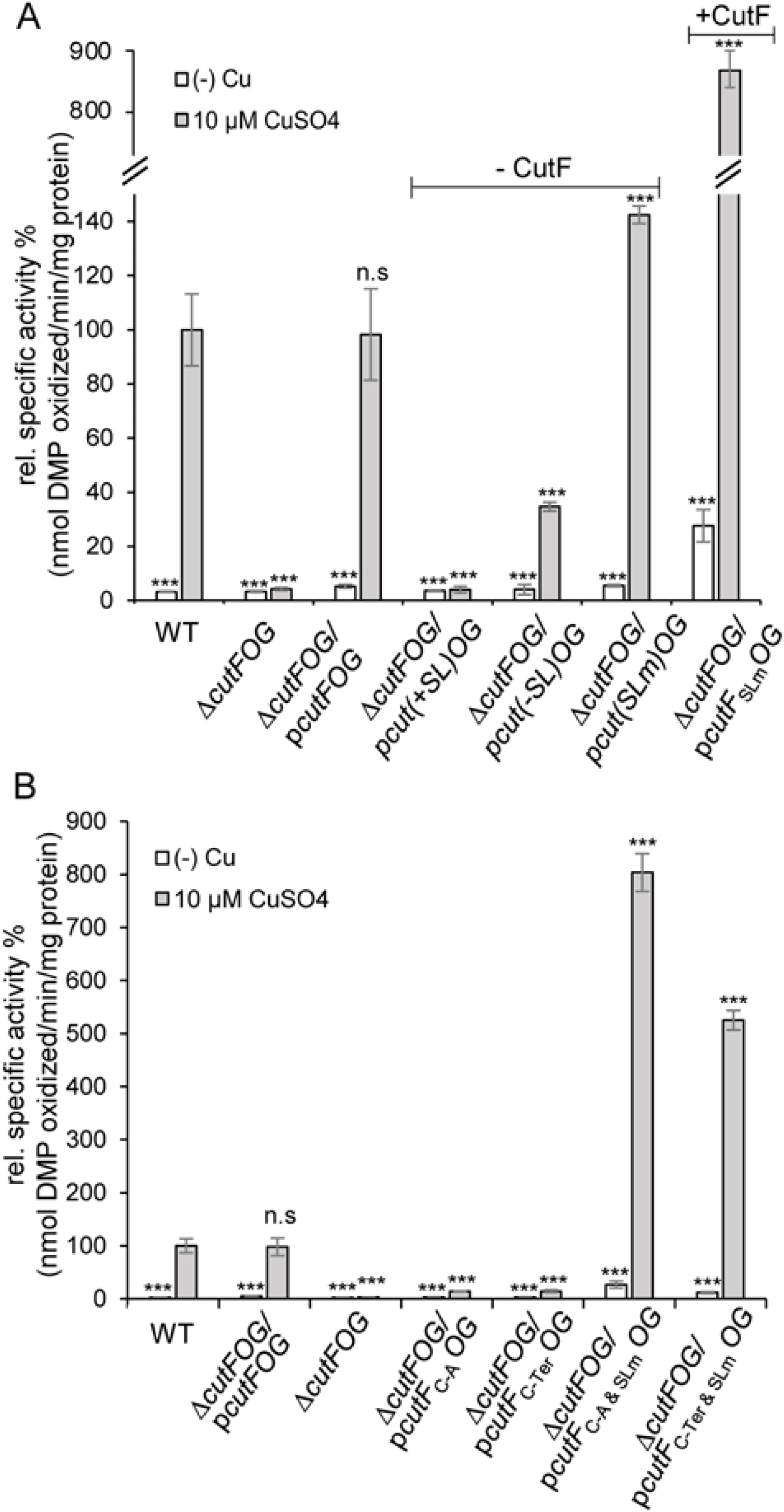
CutF is essential for Cu-induced CutO production in the presence of the stem-loop, but not in its absence. (**A**) CutO activity was determined as described in Fig. 1. p*cutF(SLm)OG* is a derivative of p*cutFOG* in which the anti-SD within the SL was mutated. p*cut(+SL)OG* encodes just *cutOG* and contains the stem-loop upstream of *cutO*, while p*cut(-SL)OG* lacks the stem-loop, and p*cut(SLm)OG* contains *cutOG* with the mutated anti-Shine-Dalgarno sequence (see **Fig. S2A** for details). The presence or absence of *cutF* is indicated on top of the figure. (**B**) CutO activity of *R. capsulatus* Δ*cutFOG* cells expressing *cutF* with a mutated Cu-binding motif (C-A) or a truncated C-terminus (ΔC-Ter) combined with an intact or mutated stem-loop (Slm). All mutations were located on the p*cutFOG* plasmid carrying the tagged versions of the genes. The CutO activities of wild-type (WT), the Δ*cutFOG* strain and the Δ*cutFOG* carrying the p*cutFOG* plasmid served as controls. The CutO activities of the indicated strains in (A) and (B) and statistical analyses (n=9) were calculated as before.

### Conserved motifs of CutF cooperate with SL for Cu-dependent CutO production

CutF contains a putative Cu-binding motif (CxxxC), commonly seen in Cu-binding proteins, and a conserved C-terminal proline-rich motif, reminiscent of ribosomal stalling sequences (26, 27) (**Fig. 1A**). Both motifs are required for Cu-dependent CutO activity (15). Whether these motifs are important for increasing the accessibility of the *cutO* SD site was tested by using strains that expressed mutated CutF variants combined with *SLm*. When the cysteine residues of the Cu-binding motif were replaced by alanine (CutF_C-A_) in the presence of wild type SL, no CutO activity was detectable. The same was also observed for a CutF variant lacking the proline-rich sequence (CutF_ΔC-Ter_). Conversely, both variants produced large amounts of Cu-dependent CutO activity when combined with *SLm* (**Fig. 2B**). Moreover, cells carrying either one of these double mutations (*cutF_C-A_* and *SLm or cutF_ΔC-Ter_* and *SLm*) produced much higher amounts of CutO than wild-type cells or Δ*cutFOG* cells complemented with a plasmid carrying *cutFOG* in absence or presence of Cu **(Figs. S2C & S2D)**. Thus, both the Cu-binding and the proline-rich motifs of CutF cooperate with SL for maximal Cu-dependent CutO production.

### The Sec signal peptide of CutF is required for Cu-dependent CutO production

A striking feature of the proteins encoded by *cutFOG* is that CutF has a predicted Sec signal peptide, whereas CutO and CutG contain Tat signal peptides that are typical for proteins translocated across the membrane in a folded, or partially folded state (28). As folding of Tat substrates occurs after their release from the ribosome, the Tat-system transports proteins post-translationally (29). In contrast, the Sec-system can act both post- and co-translationally (30, 31). Consequently, co-translational export of CutF might control the translation of the down-stream encoded *cutO*. In this case, its Sec signal peptide should be essential for controlling CutO production by coupling CutF export to *cutO* translation. Indeed, when the signal peptide of CutF was deleted (*cutF*_ΔSP_*OG*), the activity and steady-state levels of CutO were decreased to the background levels of the Δ*cutFOG* strain (**Figs. 3A** **& S3A)**. However, the signal peptide of CutF was dispensable when SL was mutated (**Figs. 3A** **& S3B**). Thus, CutF export to the periplasm is required for CutO production only in the presence of SL. Whether CutF export occurs via the SecYEG translocon was tested by replacing the Sec signal peptide of CutF with the Tat signal peptide of NosZ. This replacement resulted in a drastic decrease of CutO activity (**Fig. 3A**) and reduced levels of CutO **(Fig. S3A)**. Thus, seemingly, translocation of CutF into the periplasm *per se* is insufficient, and CutF requires its Sec signal peptide to support full CutO production.

**Figure 3.**
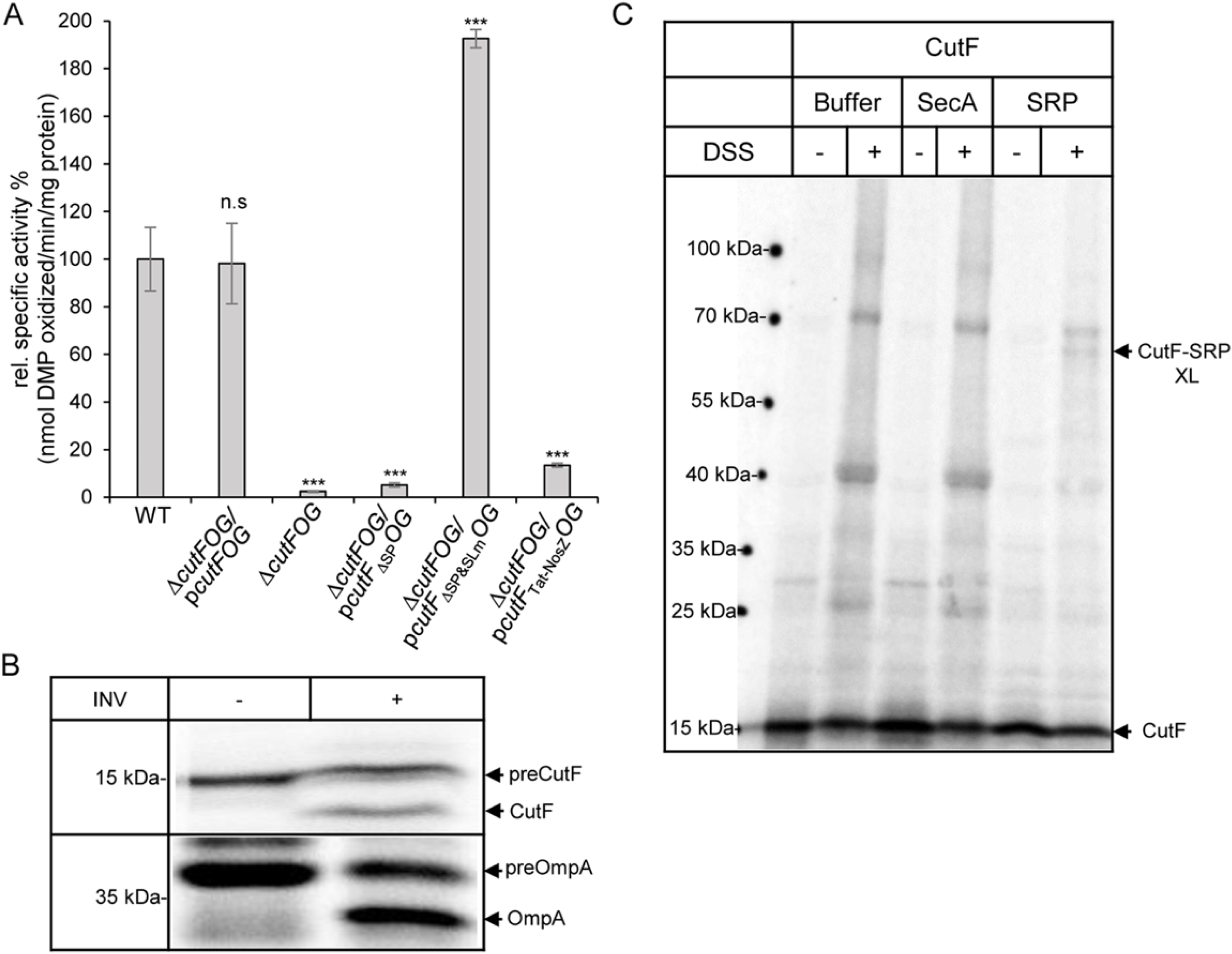
The Sec signal peptide is essential for CutF function. (**A**) CutO activity of *R. capsulatus* Δ*cutFOG* cells carrying p*cutFOG* variants with either deletion of the *cutF* signal sequence (p*cutF_Δ_*_SP_*OG*) or a replacement of its Sec signal sequence with the NosZ twin-arginine signal sequence (p*cutF_Tat-NosZ_OG*). The deletion/replacement of the Sec signal sequence was also combined with the stem-loop mutation in the plasmid p*cutF*_ΔSP&SLm_*OG*. Cells were grown on MPYE medium supplemented with 10 µM CuSO_4_. The CutO activities of the indicated strains and statistical analyses (n=9) were calculated as before. (**B**) An *E. coli* transcription/translation system was employed for *in vitro* synthesis of CutF in the absence and presence of *E. coli* inverted inner membrane vesicles (INV). The *E. coli* protein OmpA was used as a control. After *in vitro* synthesis, the radioactively labelled samples were separated by SDS-PAGE and analyzed by phosphorimaging. Indicated are the signal sequence containing precursors of CutF and OmpA (preCutF, preOmpA), and the mature CutF and OmpA proteins. (**C**) CutF was synthesized *in vitro* and incubated with either buffer or 36 ng/µl of purified SecA or signal recognition particle (SRP). Samples were subsequently treated with disuccinimidyl suberate (DSS) or buffer, separated by SDS-PAGE and analyzed by phosphorimaging. Indicated are the *in vitro* synthesized CutF and the CutF-SRP cross-linking product (CutF-SRP XL).

Although CutF has yet to be detected *in vivo*, its translation proficiency was confirmed using an *in vitro* transcription-translation system of *E. coli* (15). This system was used to determine whether the Sec signal sequence of CutF was cleaved in the presence of *E. coli* inner membrane vesicles (INVs) as a further proof of its translocation by the SecYEG translocon. When CutF was synthesized *in vitro* in the absence of INVs, a single, radioactively labelled band corresponding to the expected size of CutF was seen (**Fig. 3B**). In the presence of INVs, a second band of lower molecular weight appeared, corresponding to mature CutF lacking its signal peptide. As a control, signal sequence cleavage was monitored for OmpA, which is a model substrate used for protein translocation and signal peptide cleavage in *E. coli* (34)(**Fig. 3B**). Thus, CutF has a cleavable Sec signal peptide and can be translocated by the SecYEG translocon.

### Co-translational export of CutF and role of its C-terminal proline-rich motif

Targeting of proteins containing Sec signal peptides to the SecYEG translocon can occur via either SecA or signal recognition particle (SRP). SecA preferentially directs secretory proteins post-translationally to the SecYEG translocon, whereas SRP primarily targets membrane proteins co-translationally (30). Whether CutF interacts with SecA or SRP was probed using an *in vitro* cross-linking approach. *In vitro* synthesized CutF was incubated with purified SecA or SRP, followed by chemical cross-linking using disuccinimidyl suberate (DSS) (**Fig. 3C**). In the presence of SRP, addition of DSS resulted in a radioactively labelled band at 65 kDa, corresponding in size to a cross-link between SRP and CutF. No specific cross-linking product was detected in the presence of SecA. Although the data do not exclude that CutF might also interact with SecA, they show that SRP can target CutF to SecYEG and support co-translational export of CutF.

The C-terminus of CutF contains a conserved proline-rich sequence required for Cu-dependent CutO production that is reminiscent of ribosomal stalling sequences. (**Figs. 1A** **& 2B**) (15). Cis-acting translational modulators containing C-terminal ribosomal stalling sequences frequently execute unfolding of SL structures that cover the SD sequences of downstream genes (32, 35, 36). To probe whether the proline-rich motif of CutF acts as a ribosomal stalling sequence, *in vivo* metabolic labeling experiments were performed in *E. coli*. CutF was expressed under *T7*-promoter control, endogenous transcription was blocked by rifampicin, and cells were supplemented with ^35^S-labelled methionine/cysteine. The data show that CutF is not detectable either in the absence or presence of Cu unless its signal sequence is deleted (CutF_ΔSP_) (**Figs. 4A** **& S4A**). This indicates that the cytoplasmic form of CutF is stable, but that it is rapidly degraded upon translocation into the periplasm. This observation is consistent with a plausible role of CutF serving as a transient sensor monitoring the periplasmic Cu content. However, as neither CutF nor its variant lacking the proline-rich motif (CutF_ΔC-ter_) was detectable, it remained unanswered whether its C-terminus contains a ribosomal stalling sequence .

**Figure 4.**
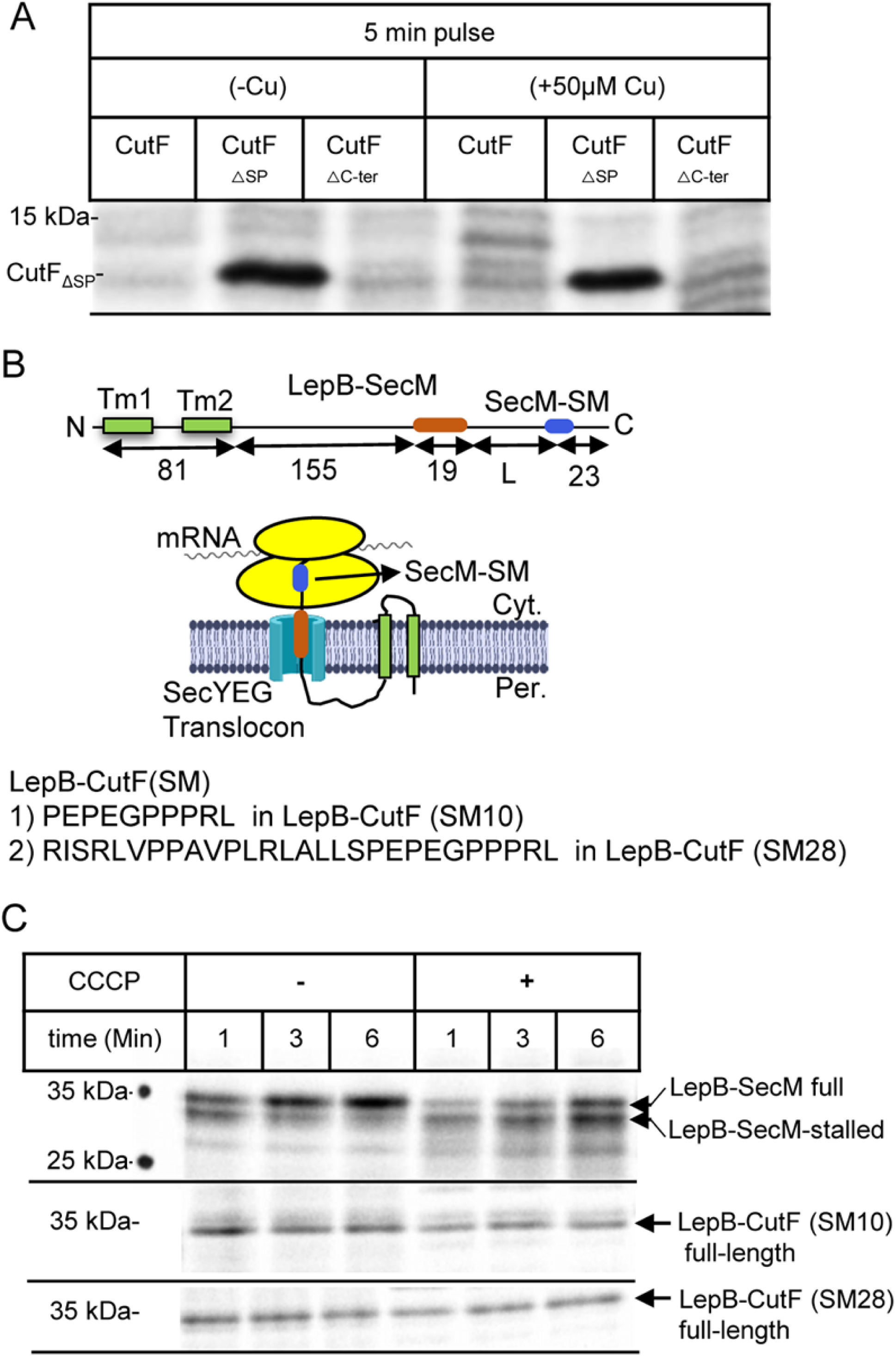
Secreted CutF is rapidly degraded. (**A**) CutF and its variants lacking either the signal sequence (CutF_ΔSP_) or the C-terminal proline-rich motif (CutF_ΔC-ter_) were expressed *in vivo* under *T7*-promoter control in *E. coli* cells and labeled with [^35^S] methionine/cysteine. After 5- and 10-min labelling, whole cells were precipitated with trichloroacetic acid (TCA), separated by SDS-PAGE and analyzed by phosphorimaging. When indicated, labeling was performed in the presence of 50 µM Cu. The complete phosphorimaging covering 10-min labelling is shown in **Fig. S4A**. (**B**) Cartoon showing the features of the translational stalling sensor LepB-SecM, which contains the first two transmembrane domains of LepB fused to a hydrophobic stretch (H) and the SecM stalling motif (SecM-SM) (upper panel). Sequences of the CutF stalling motifs that were used to replace the SecM-SM in the LepB stalling sensor, resulting in LepB-CutF(SM10) and LepB-CutF(SM28) (lower panel). (**C**) LepB–SecM, LepB-CutF(SM10) and LepB-CutF(SM28) were expressed *in vivo*, radio-labelled, and processed as in (A). When indicated, the protonophore CCCP was added. Indicated are the full-length LepB–SecM (upper band), its stalled version (lower band), and the full-length versions of LepB- CutF(SM10) and LepB-CutF(SM28).

Many ribosomal stalling sequences have been identified and characterized by fusing them to upstream sequences, such as the leader peptidase LepB (37, 38) (**Fig. 4B**). Indeed, *in vivo* pulse-labelling of LepB fused to SecM (LepB-SecM) with its intact ribosomal stalling motif (SecM-SM) results in two bands, reflecting full-length and stalled forms (**Fig. 4C**). Other studies have shown that the formation of a stalled LepB-SecM band indeed requires an intact SecM stalling motif (37). When the SecM-SM motif of LepB-SecM is replaced with the C-terminal 10 (SM10) or 28 (SM28) amino acids of CutF encompassing its proline-rich motif, resulting in LepB-CutF (SM10) or (SM28) (**Fig. 4B**), only the full-length proteins, but not the stalled forms, were detectable (**Fig. 4C**).

Over time, the arrested LepB-SecM is converted to its full-length, demonstrating that protein translocation across the membrane provides sufficient force to extract the stalling peptide out of the ribosome (37, 38). The protonophore CCCP inhibits protein translocation (37, 38) and reduces conversion of the stalled LepB-SecM into full-length (**Fig. 4C**). Similar analyses with LepB-CutF show that the addition of CCCP slightly reduced its production, but no corresponding stalled form was observed (**Fig. 4C**). Thus, the proline-rich sequence of CutF does not act *per se* as a strong stalling sequence when tested using LepB-SecM. However, as these constructs lacked the Cu-binding motif any effect of Cu could not be tested.

### Cu addition decreases co-translational export of CutF

As Cu-dependent CutO production requires both the Cu-binding and the proline-rich motifs of CutF (**Fig. 2B**) (15), the mature part of CutF (90 amino acids) was fused to the first two transmembrane domains of LepB. Further, a 23 amino acids long peptide was added to the C-terminus after the proline-rich motif, generating a 194-residue long LepB-CutF fusion to detect its possible arrested form (**Fig. 5A**). Pulse labelling resulted in a ∼21 kDa product, which corresponds to the full-length LepB-CutF as the dominant species. However, the arrested LepB-CutF, which should migrate around 19 kDa, was not detected (**Fig. 5B**, left panel). Instead, several bands of ∼12 to 14 kDa were seen, likely corresponding to proteolytic cleavage products of the secreted CutF portion of LepB-CutF. The proteolytic sensitivity of secreted CutF (**Fig. 4A**), and the stability of the LepB-SecM (37) and LepB-CutF(SM) fusions (**Fig. 4C**) were seen before.

**Figure 5.**
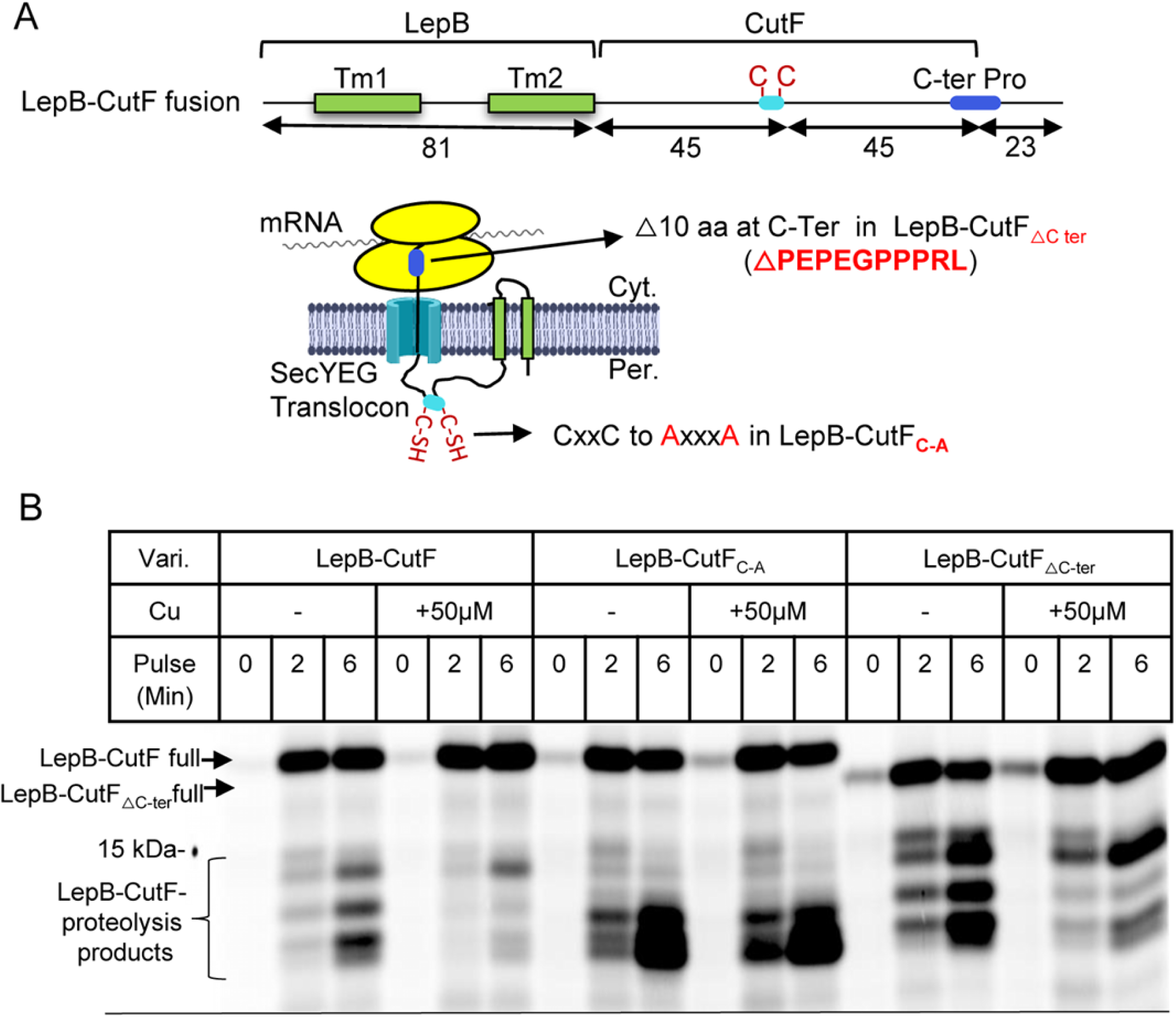
Cu enhances translational stalling and reduces proteolysis of LepB-CutF fusion. (**A**) Cartoon representation of the LepB-CutF fusion and its variants lacking either the Cu-binding motif (LepB-CutF_C-A_) or the C-terminal proline-rich motif (LepB-CutF_ΔCter_). The mature part of CutF and of both variants were fused to LepB immediately after the second TM helix. (**B**) Pulse-labeling of LepB-CutF and both variants as described in Fig. 4 in the presence or absence of 50 μM CuSO_4_. After the 0-, 2- and 6-min pulses, the samples were precipitated and analyzed by phosphorimaging. Indicated are full-length LepB-CutF and the proteolysis fragments.

Provided that this proteolysis is indeed associated with export of CutF into the periplasm, we reasoned that conditions that enhance its translational stalling and consequently decrease its export, should reduce proteolysis. When pulse labeling was repeated in the presence of Cu (50 µM), only full-length of LepB-CutF and its 14 kDa fragment were visible. The other proteolytic cleavage products were largely absent (**Fig. 5B**, left panel), suggesting that Cu addition enhanced translational stalling of LepB-CutF. However, it was recently shown that Cu can also inhibit protein export via the Sec61 translocon (39). To exclude the possibility of Cu-induced inhibition of the homologous SecYEG translocon, the LepB-SecM fusion was used as a control **(Fig. S4B**). Addition of Cu did not inhibit translocation of LepB-SecM, confirming that reduction of LepB-CutF proteolysis does not arise from inhibition of the SecYEG translocon. Thus, Cu-induced reduction of LepB-CutF proteolysis suggests its decreased export, possibly via a ribosomal stalling-like process.

This Cu-dependent process was further investigated using the LepB-CutF variants lacking either the Cu-binding or the proline-rich motifs (LepB-CutF_C-A_ or LepB-CutF_ΔC-ter_) (**Fig. 5A**). When the pulse labelling experiment was repeated with LepB-CutF_C-A_ or LepB-CutF_ΔC-ter_ (**Fig. 5B**, middle and right panels, respectively) the addition of Cu at best slightly decreased proteolysis, unlike the wild type LepB-CutF case. Thus, the occurrence of this ribosomal stalling-like process requires the presence of both Cu-binding and proline-rich motifs of CutF. The identity of the LepB-CutF proteolysis products, and the basis of the slightly different proteolysis patterns seen with different LepB-CutF variants remain unknown. A possibility is that they might arise from their slightly different structures and interactions with Cu, but this needs to be further explored.

### CutF-like proteins are clustered in several sequence similarity network clusters

Our earlier finding that CutF-like proteins are widespread in Pseudomonadota (15), combined with the insights emerging during this work, enticed us to inquire about the conservation of the sensing mechanisms by CutF-like proteins, and the identity of proteins that they may regulate. A large-scale bioinformatic approach analyzing available protein sequences, their genomic contexts, and the putative intergenic RNA motifs found downstream of *cutF-*like genes was used. Starting with our previous rule-based list (**list A,** 317 entries, **Fig. 6A**) of CutF-like proteins encoded by genes neighboring *cutO* (15), an iterative search with *jackhammer* against reference proteomes in the UniProt database was carried out. Unlike previously (15), the analysis was not limited to Pseudomonodota and sequences with matches to annotated domains were not removed. Several filters were applied for sequence characteristics at the protein (< 150 amino acid long, presence of a proline-rich (PP) motif within the 15 C-terminal residues and an N-terminal putative signal peptide (SP)), and at the intergenic (downstream gene on the same strand with an intergenic space of < 1,000 nucleotides) levels (see Materials and Methods for details) (**Fig. 6A**). This yielded an initial list (**list B**) with 2,540 proteins that can be clustered into 29 sequence similarity network (SSN) clusters (1 to 29, each with > 10 members) (**Figs. 6C** **& S5**). The hidden Markov model (HMM)-profile generated for each of the 29 SSN clusters showed distinct putative metal-binding sites with cysteines, histidines, or a combination of both (**Fig. S6**). In particular, the SSN clusters 14 and 17 comprise CutF-like proteins with only histidine-rich motifs (**Fig. S6**).

**Figure 6.**
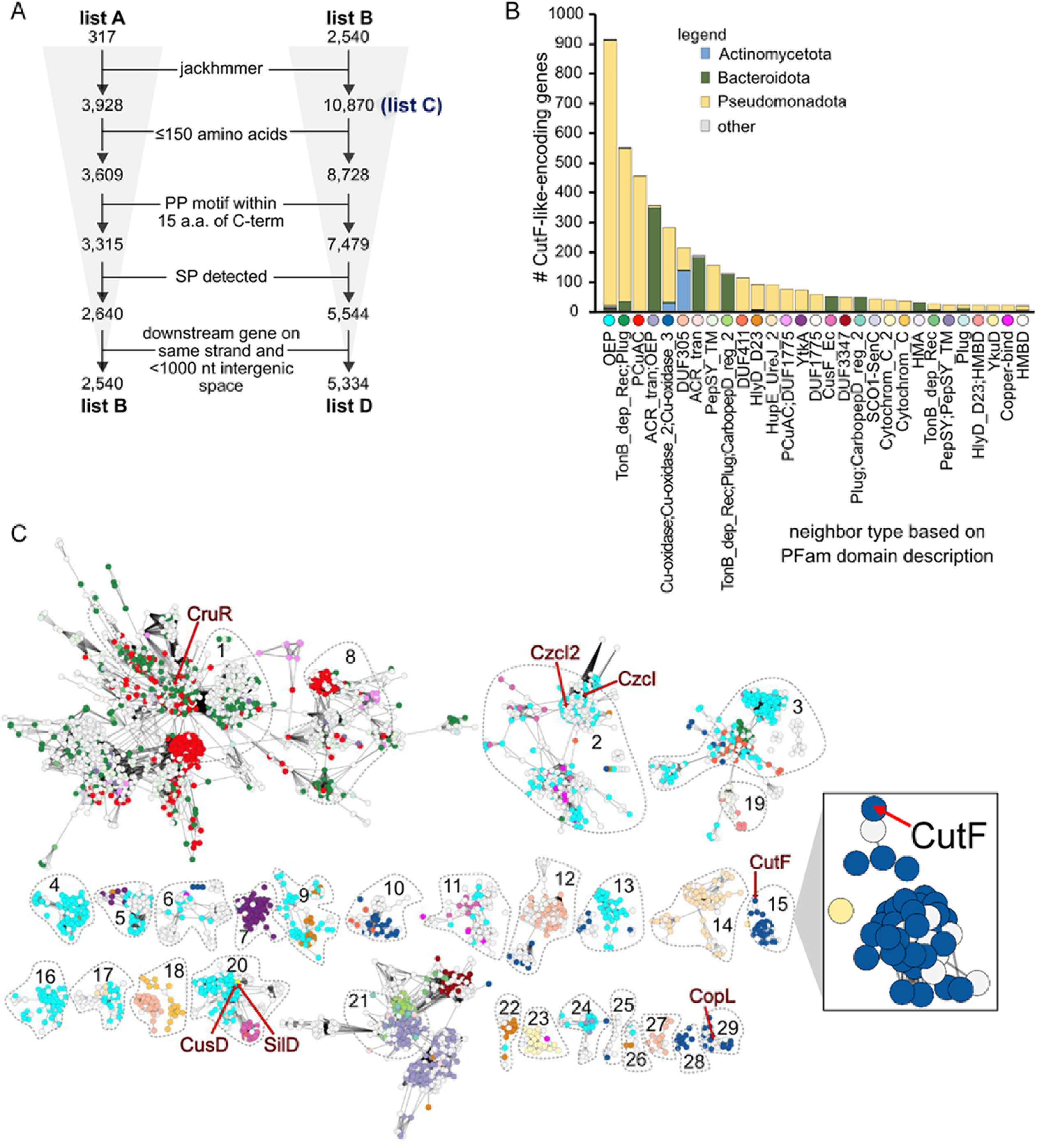
Identification and cluster analyses of CutF-like proteins. **(A)** Analysis workflow for identification of CutF-like proteins used in downstream analyses. List A is from our previous studies (15). Lists B, C, and D were generated in this study. **(B)** Types of next-door neighbors identified downstream of CutF-like genes from list D. The Pfam domain description for each class of neighbor is shown. Only neighbor classes with at least 20 instances (i.e., in 20 separate genomes) are shown. Phyla other than Pseudomonadota, Actinomycetota, and Bacteroidota are minor (i.e., no more than 5 examples for any given category; bars represented these minor “other” phyla are in grey). **(C)** The network containing proteins from list D (alignment score 5, nodes collapsed based on 100% similarity). List B protein clusters are outlined and numbered. Nodes not connected to one of these clusters are not shown but can be found in Fig. S4. Nodes representing CutF, CruR, CopL, SilD and CusD are labeled. Nodes (each circle in the network) are colored based on the downstream neighbour according to panel B. The CutF cluster is duplicated and enlarged on the right.

To increase the detection of CutF-like proteins, these 29 SSN clusters were individually searched with *jackhammer* against Reference Proteomes in the UniProt database. The results were combined in a list (**list C**, 10,870 entries), and the same filters used for List B were applied (**Fig. 6A**). The final list (**list D**) contains 5,334 CutF-like proteins from 2,804 unique UniProt proteomes. These proteins are unequally distributed among many phyla, with most of them being from Pseudomonadota (55%) and Bacteriodota (40%) (normalizing to the number of available proteomes with at least one CutF-like protein in each case; otherwise, 76% and 19%, respectively), whereas those from Chloroflexota and Spirochaetota are scarce (**Fig. S7A**).

### Genomic contexts of CutF-like proteins indicate that not all are Cu-specific, nor iso-functional

Genomic context analyses indicate that most CutF-like proteins with cysteine-rich motifs (*i.e.,* except the SSN clusters 14 and 17) are located upstream of genes involved in Cu homeostasis (**Figs. 6B) (File S1**). These include **<u>o</u>**uter membrane **<u>e</u>**fflux **<u>p</u>**roteins (OEP) associated with RND efflux systems (OEP; PF02321, often annotated as CusC/CzcC/SilC), acriflavine resistance proteins (ACR_tran; PF00873; often annotated as CusA/CzcA/SilA), CusB/CzcB (HlyD_D23; PF16576) and CusF (CusF_Ec; PF11604) (**Fig. 6B**). After OEP, the next most frequent neighbors of CutF-like proteins are TonB-like proteins and PCu_A_C-like periplasmic Cu chaperones. Strikingly, the neighboring genes to the CutF-like proteins with only histidine-rich motifs (SSN clusters 14 and 17, **Fig. S6**) belong to the HupE/UreJ family of proteins, often implicated in nickel and cobalt homeostasis. Thus, the overall findings suggest that not all CutF-like proteins are Cu-specific, and some might bind other metal ions, such as nickel and cobalt.

List D contains several previously studied proteins in addition to *R. capsulatus* CutF located in the SSN cluster 15. For example, CruR from *Bordetella pertussis* (encoded by *bp2923*), located in SSN cluster 1 (**Fig. 6C**), was recently identified as an upstream ORF (uORF) that post-transcriptionally regulates the production of a TonB-like transporter. However, in the presence of Cu, CruR was suggested to relieve ribosomal stalling (41), unlike CutF that enhances a similar process. Thus, CutF and CruR are apparently not iso-functional. Further, CopL from *Stenotrophomonas maltophilia* (encoded by *smlt2449*) located in SSN cluster 29 is the most similar protein in UniProt to CopL from *Xanthomonas perforans. Xanthomonas copL* is required for Cu-regulated expression of the downstream copper oxidase CopA, and like *cutF*, *copL* does not act *in trans*, suggesting that CutF and CopL may be iso-functional (42).

### The modes of action of different CutF-like proteins are seemingly different

Provided that *R. capsulatus* CutF achieves Cu-dependent CutO production by unfolding a downstream SL via a translational stalling-like mechanism, we inquired whether this process is also employed by other CutF-like proteins for regulating the translation of their neighboring genes. We determined the length of the intergenic regions between the *cutF*-like genes and their downstream-genes as SL unfolding by translational stalling on upstream proteins is likely limited to short intergenic regions. Determining the length of the intergenic regions downstream the CutF-like genes in List B or List D shows that most are below 100 nucleotides (**Figs. 7A** **& S7B)**. Remarkably, some CutF-like proteins have very short intergenic regions (< 10 nucleotides) or even overlap with their downstream neighbor. Nearly 60% of these cases encode either the cuproenzyme nitrous-oxide reductase, or its transcriptional regulator NosR/NirR (43). This suggests that other type(s) of co-translational Cu-sensing mechanism(s), distinct from that seen with *R. capsulatus* CutF might regulate nitrous oxide reductase production. Further, although most CutF-like proteins contain putative Sec signal sequences like *R. capsulatus* CutF (**Fig. 7B**), not all down-stream proteins have a Tat signal sequence like CutO. Indeed, the down-stream proteins frequently contain Sec and other signal sequences (**Figs. 7B** **& S7C**), further suggesting that not all CutF-like proteins employ identical modes of regulation.

**Figure 7.**
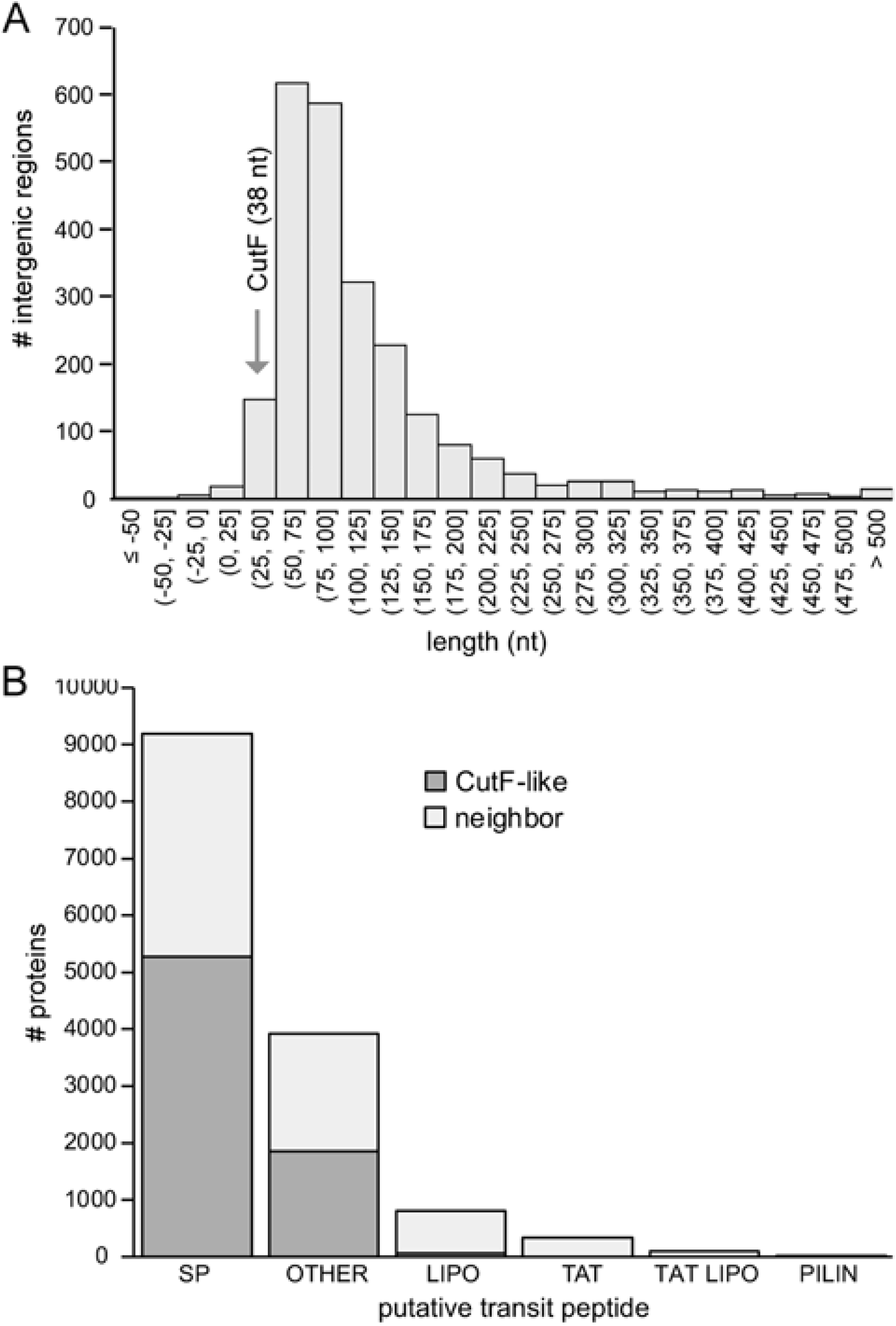
Features associated with CutF-like genes/proteins and neighbours. **(A)** Distribution of intergenic lengths between genes encoding CutF-like proteins and the downstream gene from list B. **(B)** Presence of signal sequences in CutF-like proteins and the down-stream encoded proteins, shown individually. Abbreviations: SP, Signal Peptide (Sec/SPI); LIPO, Lipoprotein signal peptide (Sec/SPII); TAT, TAT signal peptide (Tat/SPI); TATLIPO, TAT Lipoprotein signal peptide (Tat/SPII); PILIN, Pilin-like signal peptide (Sec/SPIII); Other refers to no signal peptide detected.

## Discussion

CutF belongs to the abundant DUF2946 protein family, and our bioinformatics analyses show that they are frequently found upstream of genes encoding proteins involved in heavy metal detoxification. Most CutF-like proteins contain a cleavable Sec signal sequence, a Cu-binding motif in the mature part, and a C-terminal proline-rich sequence reminiscent of translational stalling sequences. The presence of an intact *cutF* located immediately upstream of *cutO* is essential for Cu-dependent production of the multi-copper oxidase CutO, even though CutF is not detected in whole cells. This may suggest that *cutF* produces a regulatory RNA that controls CutO production. However, our data demonstrate that translation of *cutF* is required for CutO production, followed by rapid CutF proteolysis: (i) *in vitro* transcription/translation experiments confirm the production of CutF and cleavage of its signal sequence in the presence of membranes. (ii) In cells, replacing the initiator ATG codon of *cutF* with a TAG stop codon prevents CutO production. (iii) *In vivo* pulse-experiments show cytoplasmic accumulation of CutF in the absence of its signal sequence, indicating that proteolysis occurs after export into the periplasm. (iv) CutF supports full CutO production only when secreted into the periplasm via a Sec signal sequence, but not when this Sec signal sequence is deleted or even replaced by a Tat signal sequence. Moreover, Cu-dependent CutO production also requires the presence of an intact Cu-binding motif in CutF.

Our data further demonstrate that the function of CutF is invariably linked to the presence of the SL sequence between *cutF* and *CutO*. Previous data had indicated that this SL might shield the Shine-Dalgarno sequence of *cutO* and requires unfolding to allow *cutO* translation (24). Our data now show that translation and co-translational export of CutF are both required for this unfolding event in conjunction with the presence of intact Cu-binding- and proline-rich motifs. Altogether, these observations support the role of CutF as a periplasmic Cu sensor regulating *cutO* translation in response to extra-cytoplasmic Cu availability.

Proline-rich motifs often act as ribosomal stalling sequences and slow-down translation due to inefficient peptide-bond formation at the ribosome (26, 27). The presence of positively charged residues in close proximity to the proline-rich motif, as observed for CutF (**Fig. 1A**), further reduces translational speed (44). C-terminal stalling sequences that regulate translation initiation of down-stream genes have been observed with several proteins. Examples are the secreted proteins SecM, VemP or MifM, which regulate the production of SecA, SecDF and YidC2, respectively, in response to the cellular protein export status (32, 33, 35, 36). Release of SecM-, VemP- or MifM-induced translational stalling depends on the SecYEG translocon, which provides sufficient force to extract the stalling sequence out of the ribosome. When SecYEG activity declines, translational stalling leads to unfolding of a SL in the intergenic regions and increases translation initiation of the down-stream ORFs on the same mRNA (32, 33, 35, 36). SL unfolding is likely achieved by the ribosomal RNA helicase activity, which can unwind RNA secondary structures that are in its proximity (45).

SecM is co-translationally secreted by the SRP pathway and rapidly degraded in the periplasm (46), which we also observe for CutF. In the absence of Cu, CutF is secreted into the periplasm and degraded, as indicated by the observation that (i) CutF is only detectable in pulse-labelling experiments when the signal sequence is removed, and (ii) the LepB-CutF fusion construct is proteolyzed in the absence of Cu. However, unlike SecM, the ribosome stalling-like sequence of CutF *per se* does not seem sufficient to achieve its translational stalling during export via the SecYEG translocon. This is seen with the LepB-CutF (SM10 or SM29) fusions, where the SecM stalling sequence is substituted by the proline-rich (SM10 or SM29) sequences of CutF. This is also seen with the LepB-CutF fusion in the absence of Cu, despite the presence of the Cu-binding motif of CutF. However, in the presence of Cu, the translational stalling-like process that CutF undergoes apparently is prolonged, as suggested by its decreased proteolysis due to the inaccessibility of its degradation sites caused by slowed translation. This occurs only when both conserved Cu-binding and ribosome stalling-like motifs of CutF are present, as seen with the LepB-CutF_C-A_ and LepB-CutF_ΔC-ter_ variants. The Cu-dependent prolonged ribosomal stalling potentially unfolds the SL and renders the SD of *cutO* accessible for translation initiation (**Fig. 8**). Accordingly, Cu-dependent CutO production requires the presence of Cu, Cu-binding, and C-terminal proline-rich stalling-like motifs of CutF, as well as its co-translational export via a Sec signal sequence. As a corollary of this working model, in the absence of the SL, CutF and its characteristic features are dispensable, as shown here.

**Figure 8.**
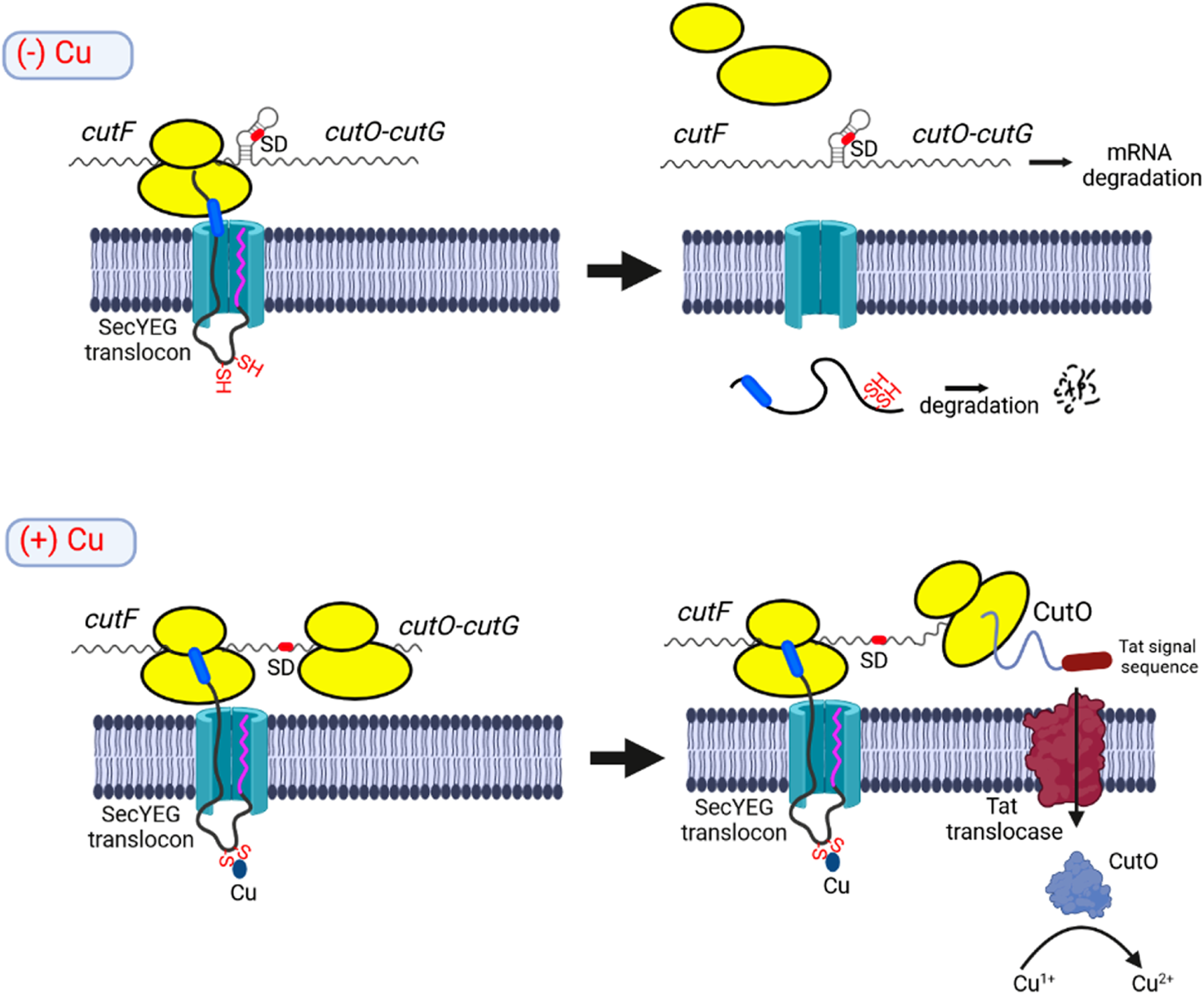
Model for Cu-controlled CutO production via the periplasmic Cu sensor CutF. CutF contains a cleavable signal sequence (magenta) and is co-translationally targeted to the SecYEG translocon by SRP. In the absence of Cu (upper panel), the C-terminal stalling sequence (blue) is pulled out of the ribosome by the translocation activity of the SecYEG translocon and CutF is completely synthesized and secreted into the periplasm, where it is rapidly degraded. Complete synthesis of CutF and ribosome dissociation prevents the unfolding of the down-stream stem-loop, which shields the Shine-Dalgarno sequence (SD; red) of *cutO*. Consequently, the *cutFOG* mRNA is degraded. In the presence of Cu (lower panel), the nascent CutF protein binds Cu via its CxxxC motif, which transiently arrests the stalling sequence within the ribosomal tunnel. This provides a time window for the helicase activity of the ribosome to unfold the stem loop and to initiate *cutO* translation via the now accessible SD. CutO is then produced and post-translationally secreted via the Tat translocase into the periplasm where it oxidizes Cu^1+^ to the less toxic Cu^2+^.

Cu-induced ribosomal stalling also explains why most CutF-like proteins have a Sec signal sequence because only a Sec signal sequence allows coupling of CutF export *to cutFOG* translation. Further, the occurrence of a Sec signal sequence in CutF and Tat signal sequences in CutO and CutG is consistent with the observed physical contact between the SecYEG translocon and the Tat translocase (38, 47).

How Cu-binding to CutF apparently prolongs the translational stalling-like process seen in this work is currently unknown. A possibility could be direct or indirect conformational changes of the nascent peptide upon binding of Cu, acting synergistically with the ribosomal stalling-like motif. Another possibility is that the folding force induced by Cu-binding to CutF’s CxxxC motif could be weaker than a disulfide-bridge formation, prolongating translational arrest (48). Alternatively, the short distance (just 45 amino acids) between the Cu-binding motif and C-terminal motif may be critical. This positions the Cu-binding motif within the periplasmic vestibule of the SecY channel (31), assuming that approx. 30 amino acids are shielded within the ribosomal tunnel (49) and approx. 17 amino acids are required to cross the bacterial membrane in an unbent conformation (50). Binding of Cu to CutF CxxxC motif located in the confined space of SecY’s periplasmic vestibule could restrict the entropic force acting on the translating polypeptide (51), resulting in reduced translocation and prolonged translational arrest. Future structural studies depicting the exact conformation of CutF trapped in the ribosomal tunnel in the presence of Cu may further elucidate this process.

C-terminal stalling sequences that regulate transcription of down-stream genes have also been observed. One example is TnaC, which regulates tryptophan metabolism in *E. coli.* In the presence of tryptophan, the stalling sequence prevents the release of TnaC from the ribosome and inhibits Rho-dependent transcription termination. Consequently, transcription and translation of the downstream encoded TnaA and TnaB continue (52). Indeed, Cu-binding to the CutF-like protein *B. pertussis* CruR (SSN cluster 1) inhibits the production of downstream encoded proteins by a ribosomal stalling-like process. Upon binding of Cu, CruR triggers Rho-dependent transcription termination, preventing the production of the downstream encoded TonB-dependent transporter (41). However, unlike *R. capsulatus* CutF (SSN cluster 15), CruR activity is Sec signal sequence independent, its C-terminal proline-rich ribosomal stalling-like motif is slightly different, and the *cruR-bfgR* intergenic region is rather long (162 nucleotides) (41). Further, the occurrence of CutF-like proteins without the strict conservation of both the Sec signal peptide and the C-terminal proline-rich ribosomal stalling-like motifs are exemplified by the *Cupriavidus metallidurans* CzcI and CzcI2 lacking a C-terminal proline-motif (SSN cluster 20). CzcI is encoded upstream of *czcCBA* RND efflux system and suggested to act as a metal-sensing regulator (53). These observations illustrate that the specific modes of action of the different CutF-like proteins (SSN clusters 1 to 29) and the ensuing regulatory responses may be very different, and that their elucidation will require further studies.

In summary, in the presence of Cu, the ribosomal stalling-like motif working together with the Cu-binding motif of CutF can delay its full-length synthesis apparently long enough for ribosomes to unfold the downstream SL structure to allow *cutO* translation. This enables cells to control CutO production in response to periplasmic Cu availability. The CutF-CutO pair serves as a fast-response to prevent Cu toxicity by allowing the production of a periplasmic multi-copper oxidase, before the transcriptional regulation via the Cu-responsive cytoplasmic transcription factors, like CueR or CopR (19, 20). The toxicity and abundance of Cu in natural environments (54, 55) likely justifies the small energetic investment of producing a low abundance and rapidly degraded CutF for boosting CutO production for palliating the toxicity of increasing Cu concentrations.

More broadly, this work shows that proteins containing C-terminal stalling sequences can sense intracellular metabolites, export defects as well as periplasmic and extracellular metabolite availability. Considering the abundance of CutF-like proteins and their close genetic association with a myriad of proteins, this sensing mechanism is likely an important cornerstone for bacterial adaptation to changing environmental conditions.

## Materials and Methods

### Bacterial strains, growth conditions and plasmid construction

Bacterial strains and plasmids used in this study are described in **Table S1**. *R. capsulatus* strains were grown under respiratory (Res) conditions on magnesium– calcium, peptone, yeast extract (MPYE) enriched medium (56), or Sistrom’s minimal medium A (Med A) (57), supplemented with kanamycin, gentamycin or tetracycline as appropriate (10, 1 or 2.5 µg per mL, respectively) at 35° C. For the arabinose-inducible genes in *R. capsulatus*, liquid media were supplemented with 0.5 % L-arabinose (L-ara) at OD_685_ of 0.5-0.6 and further grown for 6 h. *E. coli* strains were grown on a lysogeny broth (LB) medium (58), containing ampicillin, kanamycin, or tetracycline (at 100, 50, or 12.5 µg per mL, respectively) as appropriate. The minimal medium M63 (18 amino acids), including all essential amino acids with the exception of cysteine and methionine, was used for *in vivo* pulse labeling experiments (38). Ampicillin at 50 µg per mL was used for M63 medium.

### Cu-sensitivity assays on plates

The growth of *R. capsulatus* strains in the absence or presence of 400 µM CuSO_4_ was monitored by streaking on plates or using spot assays (18). For spot assays, strains were grown semi aerobically overnight to an OD_685_ of ∼ 0.9, and cell counts were determined based on OD_685_ of 1.0 = 7.5 x 10^8^ cells/mL. For each strain, 1 x 10^8^ cells were resuspended in 400 µL medium and the cell suspension was subsequently serially diluted in a 96-well plate. Dilutions ranging from 10^0^ to 10^-7^ were spotted on MPYE plates by a 48-pin replica plater. The plates were incubated under the Res conditions for approx. 2 days before scoring the data.

### Construction of p*cutF*, p*cutF*_Stp_*OG* plasmids

The p*cutFOG* plasmid (15), carrying the tagged version of *cutFOG* genes (*cutF*_N-Flag_*O*_C*-*Flag_*G*_C*-*MycHis_) and covering its 526 bp upstream (promoter) and 466 bp downstream (transcriptional terminator) parts, was used for genetic manipulations. The p*cutF* (*cutF*_N-Flag_ under the endogenous promoter) was constructed by removing the 280 bp *Srf*I fragment of p*cutFOG* covering the N-terminal 93 aa of the *cutO* gene and causing a frameshift mutation for the downstream parts of the operon. After the *Srf*I digestion and gel purification of the large fragment of p*cutFOG* (Qiagen Gel purification kit; Qiagen, Hilden, Germany), T4 DNA ligase (NEB Lab, United States) was used for self-ligation of the large fragment following the manufacturers protocol. 5 ul of the ligation reaction was used for transformation of the *E. coli* HB101 strain. The isolated plasmids from selected colonies were confirmed by sequencing. p*cutF*_Stp_*OG* plasmid, carrying the start to stop codon substitution of *cutF* on p*cutFOG,* was constructed by using the cutF(EP)-F, cutFstop&noSS-R, cutFstop-F and cutter-R primers (**(Table S2)** to amplify the fragments covering start to stop substitution mutation. After PCR amplification, amplified products were treated with *Dpn*I digestion the remove the template DNA, and purified by the Qiagen PCR purification kit (Qiagen, Hilden, Germany). Fragments containing the 20 bp overlapping sequences were assembled into the linearized (KpnI/XbaI digested) pRK415 plasmid by NEBuilder^R^ HiFi assembly cloning method (NEB Lab, United States) following the manufacturers protocol. The total amount of DNA fragments used was ∼ 0.4 - 0.5 pmoles, and the vector to insert ratio was ∼ 1:2. The samples were incubated in a thermocycler at 50 °C for 60 minutes, and 4 µL of the assembly reaction was transformed to a chemically competent *E. coli* HB101 strain. The pPara-*cut*O plasmid was constructed by using the 1F-92NOQ/Pbad-R primer pair to amplify the *araC*-Promoter_Ara_ fragment from pBAD plasmid and the cutO-F/cutO-R primer pair to amplify the *cutO*_C-Flag_ from genomic DNA of *R. capsulatus* (15).

### Construction of Stem-loop (SL) manipulated derivatives of *cut* operon without *cutF*

The constructions of p*cut(+SL)OG*, p*cut(-SL)OG*, and p*cut(SLm)OG* **(Fig. S2A)** were performed by using the cutO(+SL)G-F/cutTer-R, cutO(-SL)G-F/cutTer-R and cutO(SLm)G-F/cutTer-R primers pairs respectively **(Table S2)**. The promoter region was amplified by using the cutF(EP)-F/ cutF(EP)-R primer pairs **(Table S2)**. The amplified fragments were subjected to *Dpn*I digestion to remove the template DNA (p*cutFOG*) and purified by the Qiagen PCR purification kit (Qiagen, Hilden, Germany). Fragments were integrated into the linearized (KpnI/XbaI digested) pRK415 plasmid by NEBuilder^R^ HiFi assembly cloning method (NEB Lab, United States) as described above.

### Construction of signal peptide and stem-loop manipulated CutF derivatives

The signal peptide manipulated CutF derivatives p*cutF_Δ_*_SP_*OG* and p*cutF*_Tat-NosZ_*OG* **(Table S1)** were constructed by using the primer pairs; cutF(EP)-F/cutFstop&noSS-R and cutFnoSS-F/cutFter-R for p*cutF_Δ_*_SP_*OG*, and cutF(EP)-F/cutF-Tat-R and cutF-Tat-F/cutFter-R for p*cutF*_Tat-NosZ_*OG* **(Table S2)**. Amplified fragments carrying the desired modifications were cloned to linearized pRK415 plasmid as described above. The stem-loop mutation (SLm; cttc to aaaa mutation of a-SD in stem-loop) was introduced to p*cutFOG*, p*cutF*_C-A_*OG,* p*cutF_Δ_*_C-Ter_*OG,* p*cutF_Δ_*_SP_*OG* by using the cutO-SL-F/cutO-SL-R primer pair **(Table S2)**.

The correct constructs on plasmid pRK415 were conjugated into the corresponding *R. capsulatus* strains (*ΔcutF* for p*cutF* and *ΔcutFOG* for the other constructed plasmids) via triparental conjugation (59) (**Table S1**).

### Cloning of CutF and LepB-CutF variants to T7 based expression plasmid pRS1 for *in vivo* pulse labeling, *in vitro* expression, and crosslinking experiments

The plasmid pRS1-CutF used for *in vitro* and *in vivo* experiments was described previously (15). For the construction of signal peptide and C-terminal deletion versions of CutF in pRS1 (pRS-CutF_ΔSP_ and pRS-CutF_ΔC-ter_ respectively), the Q5 mutagenesis Kit (NEB Lab, MA) was used with the mutagenic primer pairs pRScutF(noSS)-F/pRS1-R for pRS-CutF_ΔSP_ and pRS1-F/pRScutF(delC-ter)-R for pRS-CutF_ΔC-ter_ **(Table S2)**. The pRSLepB–CutF(SM10) and pRSLepB–CutF(SM28) were constructed by using the Q5 mutagenesis Kit (NEB Lab, MA) with cutF(AP 10&28)-F/ cutF(AP-10aa)-R and cutF(AP 10&28)-F/cutF(AP-28aa)-R mutagenic primer pairs carrying the 10 and 28 aa C-terminal proline-rich segment of CutF respectively **(Fig. 4B, Table S2)**. 10 ng of pRS1-LepB–SecM(Ms) (38) was used as a template and manufacturer protocol was followed. 5 µL of the KLD reaction mixture was transformed to chemically competent *E. coli* NEB® 5-alpha strain selected for Amp^r^ colonies. The resulted plasmids carrying new constructs were confirmed by DNA sequencing and transformed to MC4100 by the TSB transformation procedure (60). For the LepB-CutF fusion construct, pRSLepB– CutF(SM10) was linearized by using the cutF(AP10&28)-F/LepB-cutF-V-R primer pairs by inverse PCR to keep the first and second TM helix and the last 23 aa of lepB excluding the rest of the sequence. The *cutF*, excluding signal peptide and including a 20 bp overlapping sequence of *lepB* 2^nd^ TM helix at 5’ end and last 23 aa seq at 3’ end, was amplified from p*cutFOG* by using the primers cutF(LepB)-F and cutF(LepB)-R (**Table S2).** The PCR products were digested with DpnI to remove the template DNA used for the PCR and then were purified by a Qiagen PCR purification kit (Qiagen, Hilden, Germany). The NEBuilder^R^ HiFi assembly cloning method (NEB Lab, United States) and transformation procedures similar to those described above were used for this construction. The resulting plasmid pRSLepB-CutF(fusion) was confirmed by DNA sequencing and transformed to MC4100 by TSB transformation procedure (60). The LepB-CutF_C-A_ and LepB-CutF_ΔC-ter_ versions were constructed by using the Q5 mutagenesis Kit (NEB Lab, MA) with cutF(CtoA)-F/ cutF(CtoA)-R and lepB-cutF(dC-ter)-F/ lepB-cutF(dC-ter)-R mutagenic primer pairs carrying the CxxxC to AxxxA and Δ10 amino acids from C-terminal proline-rich segment respectively (**Table S2**). 10 ng of pRS-LepB–CutF was used as a template and manufacturer protocol was followed. 5 µL of the KLD reaction mixture was transformed to chemically competent *E. coli* NEB® 5-alpha strain selected for Amp^R^ colonies. The resulting plasmids were confirmed by DNA sequencing and transformed to MC4100 by TSB transformation procedure (60).

### Preparation of the periplasmic fraction for multicopper oxidase activity assays and SDS-PAGE

The periplasmic fraction from *R. capsulatus* cells was isolated from 50 mL overnight (∼20 h) cultures, grown on MPYE medium under Res condition at 35° C with 110 rpm shaking in the absence or presence of 10 µM CuSO_4_ (61, 62). Cells were harvested and washed at 4° C with 12 mL of 50 mM Tris-HC1 pH 8.0. The pellet was resuspended to a concentration of 10 mL/g of wet weight in SET buffer (0.5 M sucrose, 1.3 mM EDTA, 50 mM Tris-HC1 pH 8.0), and incubated with 600 µg lysozyme/mL at 30° C for 60 min. Spheroplasts formation was monitored by microscopy. Spheroplasts were collected by centrifugation (13.000 rpm) at 4° C for 30 min. The supernatant (periplasmic fraction) was directly used for multi-copper oxidase activity assays and SDS-PAGE or stored at -80° C.

The oxidation of 2,6-dimethoxyphenol (2,6-DMP) by the periplasmic fraction (50 µg total protein) was monitored as endpoint assay at 468 nm. The molar concentration of oxidized 2,6-DMP was calculated using ɛ = 14,800 M^−1^ cm^−1^ (63).

### Immune detection

Following the SDS-PAGE, proteins were electro-blotted onto nitrocellulose (GE Healthcare, Germany) or PVDF Immobilon-P (GE Healthcare, Germany) membranes, respectively and antibodies against the Flag-tags was purchased from either Sigma or Millipore (Temecula, USA).

### *In vitro* synthesis and crosslinking of CutF

*In vitro* protein synthesis was performed in an *E. coli in vitro* transcription/translation system as described before (34). Samples were incubated at 37° C for 30 min, with gentle shaking. The reaction was stopped with 5% trichloroacetic acid (TCA) for 30 min on ice. Precipitated proteins were pelleted by centrifugation (13.000 rpm) and resuspended in 30 µl TCA loading dye (64). Samples were separated on a 5-15% SDS-PAGE and visualized by phosphor-imaging.

For cross-linking of *in vitro* synthesized CutF, triethanolamine acetate pH 7.5 was replaced with HEPES/NaOH pH 7.5 buffer in the *in vitro* reaction. 36 ng/µl of purified SRP or SecA, both in 50 mM HEPES/NaOH pH 7.5, 100 mM K-acetate, 10 mM Mg acetate, 1 mM DTT, were added to the *in vitro* reaction and incubated at 37° C for 30 min with gentle shaking. SRP was reconstituted from purified Ffh and 4.5S RNA (64) and SecA was purified as described (65). Subsequently, 7.5 µl 25 mM DSS (Thermo Fischer, Germany) dissolved in dimethyl sulfoxide was added to each reaction. The samples were incubated at 25° C for 30 min and then quenched with 50 mM Tris/HCl pH 7.5. Subsequently, proteins were TCA precipitated and visualized as described above.

### *In vivo* pulse labeling

*E. coli* MC4100 strain carrying appropriate plasmids were grown at 37 °C overnight in a 3 ml LB medium containing 100 µg/ml ampicillin. Cultures (2 ml) were harvested and resuspended after two washing steps in 200 µl of M63 medium (18-aa). 150 µl of the cell suspensions were used for inoculation of 10 ml fresh M63 medium supplemented with 25 µg/ml ampicillin. The cultures were grown at 37 °C with 180 rpm shaking until OD_600_ of 0.5–0.8. Subsequently, 2×10^8^ cells were collected and transferred to 2-ml Eppendorf tubes and the volume was adjusted to 2 ml with fresh M63 medium. 50 µg/mg rifampicin was added followed by incubation for 15 min at 37 °C. The production of LepB–SecM and LepB–CutF variants was induced by the simultaneous addition of 0.1 M IPTG and 2 µl of L-[^35^S] methionine-cysteine (7 mCi/ ml, PerkinElmer Life Sciences) (38). CCCP (0.1 mM) was added when indicated, and the cultures were preincubated for 10 min at 37 °C prior to the addition of IPTG and L-[^35^S] methionine-cysteine. CuSO_4_ was added together with IPTG and L-[^35^S] methionine-cysteine. Subsequently, 100 µl of each sample was collected after different time points and directly precipitated by the addition of 10% TCA and incubation on ice for 30 min. The precipitated proteins were pelleted by 15 min centrifugation at 13,500 rpm at 4 °C. Pellets were denatured in 25 µl SDS-loading dye at 56 °C for 15 min with 1,400 rpm continuous shaking. Samples were separated by SDS-PAGE (15%) and analyzed by phosphor-imaging (38).

### Bioinformatic analyses

To generate a list of proteins that share characteristics with CutF, we used a previously generated list of CutF-like proteins, which relied on a set of rules (i.e., searched for small ORFs within a 10 gene window of the gene encoding a CutO-like protein (defined as containing PF00394 or PF07731 or both), encoded protein is smaller than 170 aa, does not match to an annotated Pfam domain, contains a signal peptide, contains CxxxC, contains PP motif, limited to Proteobacteria) (15). A multiple sequence alignment of the proteins from this list was generated with MUSCLE (66) and used to search against the UniProt database (Reference Proteomes) with jackhammer (using 6 iterations) (67). This list, named list B in **Fig. 6**, was filtered based on length (proteins longer than 150 amino acids removed), presence of a PP motif within 15 amino acids of the C-term, presence of a signal peptidase I-cleaved Sec signal peptide (as predicted by SignalP) (68), whether the neighboring downstream gene is on the same strand, and if the intergenic region with that neighboring gene is less than 1000 nucleotides. A sequence similarity network was built with these sequences using an E-value threshold of 1E-5 *(i.e.*, alignment score of 5) using the EFI-EST webtool (69). The network was visualized with Cytoscape v3.5.0 (70) and the yFile Organic layout. Clusters containing more than 10 nodes were then used to start separate jackhammer searches (i.e., the sequences in each cluster were aligned and used as a query for a jackhammer search against UniProt (Reference Proteomes) with 5 iterations). The proteins resulting from each jackhammer search were combined (resulting in list C) and filtered using the same filters as for list B (except that putative lipoprotein signal peptides transported by the Sec translocon and cleaved by Signal Peptidase II, which is detected for CruR, were included in addition to “standard” secretory signal peptides) creating list D. Genomic context information was collected using the EFI-GNT webtool (71).

## Supplemental Materials

Supplemental material for this article may be found at https://

**Figure S1. Cu-sensitivity phenotypes of different *R. capsulatus* strains.** (**A**) Cu sensitivity phenotypes of plasmid-encoded N-terminally Flag-tagged CutF variant (p*cutF*) in Δ*cutF* strain. Growth phenotypes were tested under the respiratory (Res) growth condition on MPYE medium supplemented without CuSO_4_ or with 400 µM CuSO_4_ at 35°C for 3-4 days. (**B**) Cu sensitivity of Δ*cutFOG*/p*cutF*_Stp_*OG* strain carrying the start-to-stop codon substitution of CutF was tested by spot assay. The strains were serially diluted (see the material method section) and grown as in (A). (**C**) Immunoblot analysis of the periplasmic fractions (50 mg protein) of ΔcutFOG strain carrying pPara-cutO using α-Flag antibodies. As a control, the periplasmic fraction of the ΔcutFOG strain carrying pcutFOG under the endogenous promoter was analyzed. Shown is a representative western blot of 3 biological replicates. The lower part of the SDS-PAGE was Coomassie stained and serves as a loading control. (**D**) Cu sensitivity phenotypes of stem-loop (SL) mutated derivatives of the *cut* operon lacking *cutF.* Growth conditions were as in (**A**) and details on the stem-loop mutations are depicted in Fig. S2.

**Figure S2. Construction of Stem-loop (SL) mutated derivatives of *cut* operon without *cutF.* (A)** The entire *cutF* excluding the last 4 a.a sequence was deleted from p*cutFOG* plasmid to obtain the p*cut(+SL)OG* construct. In the p*cut(-SL)OG* plasmid, the entire *cutF* and anti-SD (a-SD) sequences were deleted. The *Cut(SLm)OG* construct was obtained by introducing the CTTC-AAAA substitution mutation (SLm) (red underlined) on the a-SD sequence of p*cut(+SL)OG* plasmid to prevent stem-loop formation. SD and a-SD sequences were indicated by bold underlining letters. (**B**) Immunoblot analysis of the periplasmic fractions (100 mg total protein) of the indicated strains. α-Flag antibodies were used for the detection of CutO. A section of the corresponding SDS PAGE was stained with Ponceau-red and served as a loading control. Shown are representative western blots of three biological replicates. (**C**) Immunoblot analysis using 100 mg of protein of the periplasmic fractions of appropriate cells related to *cutF_C-A_OG* and *cutF_DC-Ter_OG* (D). α-Flag antibodies were used for the detection of CutO. Appropriate sections of the corresponding SDS PAGE or membrane blots were stained as a loading control with Ponceau-Red or Coomassie, respectively. Shown are representative western blots of three biological replicates.

**Figure S3. Immunoblot analysis of the periplasmic fractions (100 mg protein) of *R. capsulatus strains.* (A**) Δ*cutFOG* cells carrying p*cutFOG* variants p*cutF_Δ_*_SP_*OG*) and p*cutF_Tat-NosZ_OG*). (**B**) Δ*cutFOG* cells carrying p*cutF_Δ_*_SP_*OG*) and p*cutF*_ΔSP&SL_*OG* . α-Flag antibodies were used for detecting CutO. Section of the corresponding SDS PAGE was stained with Coomassie and served as a loading control. Shown are representative western blots of three biological replicates. In (**A**) and (**B**), non-relevant parts of the blot were cut out and the two parts were re-assembled, as indicated by the vertical lines.

**Figure S4. Expression of CutF, CutF_ΔSP_ and CutF_ΔC-ter_ *in vivo* under *T7*-promoter control in *E. coli* cells. (A)** After 5- and 10-min labelling with [^35^S] methionine/cysteine, whole cells were precipitated with trichloroacetic acid (TCA), separated by SDS-PAGE and analyzed by phosphorimaging. (**B**) Pulse-labeling of LepB-SecM in the presence and absence of Cu. The experiment was performed as described in Fig. 4 and full-length LepB–SecM (upper band) and its stalled version (lower band) are indicated.

**Figure S5. Analysis of the network containing proteins from list B (alignment score 5, nodes collapsed based on 100% similarity).** (A) Protein clusters containing 10 or more members are colored and labeled. Nodes representing CutF, CopL, SilD and CusD are labeled. Triangles with red outlines are from list A. (B) Nodes are colored based on the downstream gene.

**Figure S6. Sequence logos representing profile-HMM generated with list B clusters.** The number to the right refers to the cluster number in Figure 9. The “CutF” logo represents the profile-HMM from a search using CutF against UniProt using phmmer followed by jackhammer for 5 iterations.

**Figure S7. Phyla containing a CutF-like protein (A), Distribution of intergenic lengths between genes encoding CutF-like proteins and the downstream gene from list D (B), and presence of signal sequences in CutF-like proteins and the down-stream encoded proteins (C) (shown as pairs).** Abbreviations: SP, Signal Peptide (Sec/SPI); LIPO, Lipoprotein signal peptide (Sec/SPII); TAT, TAT signal peptide (Tat/SPI); TATLIPO, TAT Lipoprotein signal peptide (Tat/SPII); PILIN, Pilin-like signal peptide (Sec/SPIII); Other refers to no signal peptide detected.

**Table S1.** Strains and plasmids used in this work.

**Table S2.** Primers used in this work (5’ to 3’ direction).

## Acknowledgments

This work was supported by grants of the Deutsche Forschungsgemeinschaft to HGK (KO2184/8, KO2184/9 (SPP2002); SFB1381, Project-ID 403222702, and RTG 2202, Project-ID 278002225). CBH was supported by the US Department of Energy, Office of Science, Office of Biological and Environmental Research Awards (DE-SC0018301) and FD by the Division of Chemical Sciences, Geosciences and Biosciences, Office of Basic Energy Sciences of Department of Energy (DE-FG02-91ER20052). YÖ acknowledges support by the Philipp Schwartz Initiative of the Alexander von Humboldt Foundation and by the RTG 2202.

## Author Contributions

Conceptualization: HGK, CBH, FD; Investigation: YÖ, AA, CBH, ND, SRM; Visualization: YÖ, CBH, SRM, FD, HGK; Funding acquisition: CBH, FD, HGK; Supervision: HGK, FD; Writing: YÖ, CBH, FD, HGK. All authors have read the manuscript.

